# Strong positive biodiversity–productivity relationships in a subtropical forest experiment

**DOI:** 10.1101/206722

**Authors:** Yuanyuan Huang, Yuxin Chen, Nadia Castro-Izaguirre, Martin Baruffol, Matteo Brezzi, Anne Lang, Ying Li, Werner Härdtle, Goddert von Oheimb, Xuefei Yang, Kequan Pei, Sabine Both, Xiaojuan Liu, Bo Yang, David Eichenberg, Thorsten Assmann, Jürgen Bauhus, Thorsten Behrens, Francois Busçot, Xiao-Yong Chen, Douglas Chesters, Bing-Yang Ding, Walter Durka, Alexandra Erfmeier, Jingyun Fang, Markus Fischer, Liang-Dong Guo, Dali Guo, Jessica L.M. Gutknecht, Jin-Sheng He, Chun-Ling He, Andy Hector, Lydia Hönig, Ren-Yong Hu, Alexandra-Maria Klein, Peter Kuehn, Yu Liang, Stefan Michalski, Michael Scherer-Lorenzen, Karsten Schmidt, Thomas Scholten, Andreas Schuldt, Xuezheng Shi, Man-Zhi Tan, Zhiyao Tang, Stefan Trogisch, Zhengwen Wang, Erik Welk, Christian Wirth, Tesfaye Wubet, Wenhua Xiang, Jiye Yan, Mingjian Yu, Xiao-Dong Yu, Jiayong Zhang, Shouren Zhang, Naili Zhang, Hong-Zhang Zhou, Chao-Dong Zhu, Li Zhu, Helge Bruelheide, Keping Ma, Pascal A. Niklaus, Bernhard Schmid

## Abstract

Forest ecosystems contribute substantially to global terrestrial primary productivity and climate regulation, but, in contrast to grasslands, experimental evidence for a positive biodiversity-productivity relationship in highly diverse forests is still lacking^1^. Here, we provide such evidence from a large forest biodiversity experiment with a novel design^2^ in subtropical China. Productivity (stand-level tree basal area, aboveground volume and carbon and their annual increment) increased linearly with the logarithm of tree species richness. Additive partitioning^3^ showed that increasing positive complementarity effects combined with weakening negative selection effects caused a strengthening of the relationship over time. In 2-species mixed stands, complementary effects increased with functional distance and selection effects with vertical crown dissimilarity between species. Understorey shrubs reduced stand-level tree productivity, but this effect of competition was attenuated by shrub species richness, indicating that a diverse understorey may facilitate overall ecosystem functioning. Identical biodiversity-productivity relationships were found in plots of different size, suggesting that extrapolation to larger scales is possible. Our results highlight the potential of multi-species afforestation strategies to simultaneously contribute to mitigation of climate change and biodiversity restoration.

---

Forest ecosystems harbor around two thirds of all terrestrial plant species, but currently lose biodiversity at high rates which may threaten the production of timber, fiber, fuel and other services beneficial to humans^4^. Observational studies suggest that species-rich forests exceed the productivity of less diverse forests^5,6^, but co-varying factors (e.g. spatial heterogeneity in abiotic environment, species composition and successional stages; interventions by forest management) make assigning causation difficult. Systematic experimental manipulations of plant species composition in grassland communities^7–9^ have demonstrated that plant diversity promotes community productivity. This effect has been attributed to positive effects of niche partitioning between species, specifically to complementarity in the use of abiotic resources^10^ or interactions with enemies^11^, or to an increasing contribution of highly productive species in more diverse communities^12^. These two types of mechanisms have been related to statistical complementarity and selection effects obtained by additive partitioning^3^. However, these mechanisms may differ in species-rich forests in which neutral processes may be important^13,14^ and where “diffuse” coevolution may result in niche convergence toward generalist strategies^15^. Furthermore, trees have large and persistent vertical structures that support the long-term accumulation of biomass. Several forest experiments have recently been initiated^16,17^, but these are mainly in the temperate zone or implemented in small plots with a limited species richness gradients^18–23^. To close these critical gaps in knowledge^1^, controlled experiments in which the diversity of tree species is systematically manipulated are needed. The largest such study concerning numbers of treatments and plots has been established in 2009/2010 in subtropical south-east China and is referred to as the BEF-China experiment^2^.

Here, we report how stand-level productivity in the BEF-China experiment 3-7 years after planting was related to species richness and how variation within species-richness levels was related to trait differences among species. Experimental forest communities were constructed systematically from a pool of 40 tree (Extended Data Table 1) and 20 shrub species, and were established in plots at two hilly sites (in 2009 at site A and in 2010 at site B). By the time of our later measurements the tree communities were well established with some canopies exceeding 12 m in height in 2016. The design of previous biodiversity experiments had been criticised because not all species were found at all diversity levels, and because the compositions of the experimental communities that were realized were not nested as would be expected with sequential extinction^24^. We adopted a novel design that avoided these caveats^2^ (see Methods, Extended Data Fig. 1, Extended Data Table 2). In brief, we first created three pools of 16 species per site. These were then repeatedly split into halves, resulting in nested, non-overlapping subsets of 8, 4, 2 and 1 species. We used these sets, and in addition also the full sets of 24 species per site, to plant tree communities comprising 1 to 24 species. We further established plots with two sizes: 0.067 ha (equivalent to the Chinese area unit of 1 mu; 400 individual trees) and 0.267 ha (4 mu; 1600 individuals). The larger plots were established for one of the three 16-species pools at each site and included a split-plot treatment that consisted of understorey shrubs planted in the centre of the quadrats formed by four neighbouring trees. Shrubs were planted at a species richness of 0 (no shrubs), 2, 4 or 8, in factorial combination with the tree species-richness treatment. We assessed stand-level tree productivity in all 1-mu plots (including all 1-mu subplots of the larger plots) non-destructively by measuring stem basal area and height of the 16 central trees every year from 2013-2016 in September/October. We used these data, together with data from separately harvested trees to obtain conversion factors, to calculate tree volume and aggregated the individual volume data of live trees to the stand level. To characterize annual stand growth, we further derived yearly increments of stand volume from successive inventories. Using the same method, we determined the same metrics at the population-level (stand-level data separated into species).

**Figure 1.**
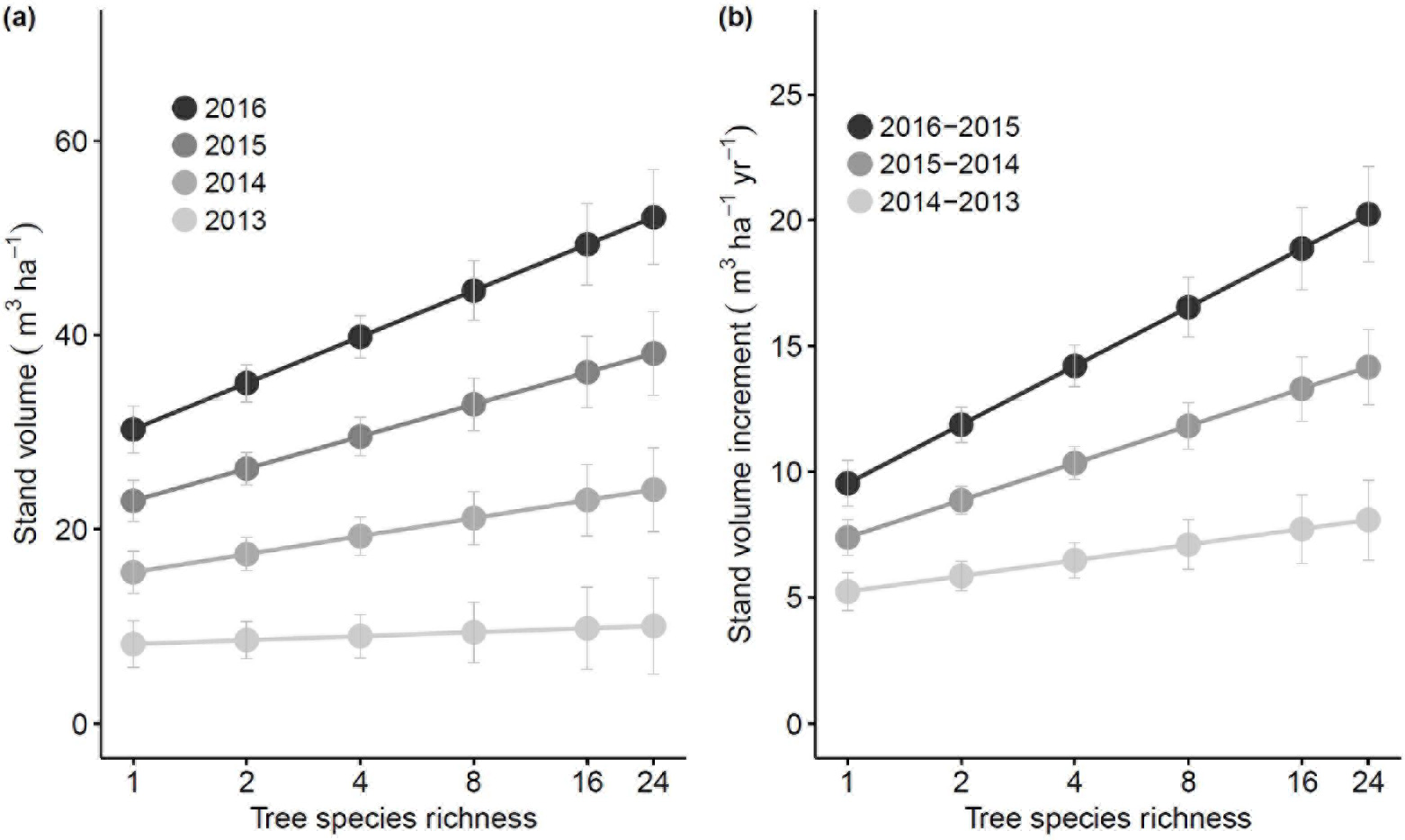
Stand-level tree volume (a), and its annual increment (b) as a function of tree species richness from 2013–2016. The figure shows predicted means and standard errors based on fitted mixed models (Table 1). Effects of species richness were significantly positive and increased significantly throughout the observation period.

**Table 1.** Mixed-effects models for effects of site, tree species richness (logSR), time (year) and interactions on stand-level tree basal area, stand-level tree volume and their increments.

| Source of variation | Basal area (n = 387) |  |  |  | Volume (n = 387) |  |  |  |
| --- | --- | --- | --- | --- | --- | --- | --- | --- |
|  | df | ddf | *F* | *P* | df | ddf | *F* | *P* |
| Site | 1 | 120.0 | 13.80 | <0.001 | 1 | 100.8 | 19.21 | <0.001 |
| LogSR | 1 | 111.2 | 6.00 | 0.016 | 1 | 88.7 | 5.26 | 0.024 |
| Year | 1 | 125.5 | 359.90 | <0.001 | 1 | 103.0 | 206.30 | <0.001 |
| Site × year | 1 | 125.3 | 5.80 | 0.018 | 1 | 103.7 | 18.80 | <0.001 |
| LogSR × year | 1 | 117.8 | 16.00 | <0.001 | 1 | 94.8 | 10.83 | 0.001 |
| Source of variation | Basal area increment (n = 387) |  |  |  | Volume increment (n = 387) |  |  |  |
|  | df | ddf | *F* | *P* | df | ddf | *F* | *P* |
| Site | 1 | 123.5 | 5.10 | 0.025 | 1 | 104.0 | 14.54 | <0.001 |
| LogSR | 1 | 115.7 | 13.30 | <0.001 | 1 | 93.9 | 9.34 | 0.003 |
| Year | 1 | 111.4 | 20.90 | <0.001 | 1 | 104.2 | 68.85 | <0.001 |
| Site × year | 1 | 115.2 | 4.40 | 0.037 | 1 | 109.2 | 27.50 | <0.001 |
| LogSR × year | 1 | 102.1 | 10.20 | 0.002 | 1 | 95.1 | 12.01 | <0.001 |

Notes:

Notes:

Fixed effects were fitted sequentially (type-I sum of squares) as indicated in the table (random terms were community composition, plot, subplot and the interaction of these with year, with site-specific variance components for species composition and plot). Abbreviations: n = numbers of plots in analysis; df = nominator degree of freedom; ddf = denominator degree of freedom; logSR = log_2_(tree species richness). *F* and *P* indicate F-ratios and the P-value of the significance test.

We found significantly positive effects of the logarithm of tree species richness on both stand volume and annual stand volume increment of trees (F_1,89_ = 5.26, P = 0.024 and F_1,94_ = 9.34, P = 0.003, respectively; Fig. 1 and Extended Data Fig. 2, Table 1). The size of these effects increased over time (F_1,95_ = 10.83, P = 0.001 and F_1,95_ = 12.01, P < 0.001, respectively, for interaction species richness × year). Similar results were obtained for stand basal area and its increment (Extended Data Fig. 3, Table 1). In 2016, at the end of our measuring period, stand basal area increased on average by 1.65 m^2^ ha^−1^ and stand volume by 5.09 m^3^ ha^−1^ with each doubling of tree species richness. After seven years of growth, the average 16-species mixture stored 22.0 ± 4.5 Mg C ha^−1^ above ground, which is double the amount found in monocultures (9.4 ± 1.1 Mg ha^−1^, Extended Data Fig. 4) and similar to the productivity of monocultures of commercial plantation species *Cunninghamia lanceolata* (22.4 ± 10.7 Mg C ha^−1^) and *Pinus massoniana* (21.0 ± 3.0 Mg C ha^−1^) that we had planted for reference at the same site (Extended Data Fig. 4, Extended Data Table 4). System-level C sequestration likely is higher, given that additional C will have been allocated to belowground tree organs^25^ and in part transferred to persistent soil pools important for long-term carbon sequestration. These strong positive effects of tree species richness were driven by faster growth of live trees in more diverse stands, and were unrelated to tree survival rate, which was independent of species richness; if anything, there was a trend towards lower survival at higher richness (Extended Data Fig. 5).

The net biodiversity effect^26^ on productivity increased through time for mixtures of all species-richness levels (Fig. 2, F_1,48_ = 23.61, P < 0.001). The positive effects of tree species richness on productivity were also reflected in a higher frequency of mixtures that overyielded relative to the ones that underyielded and in many cases of transgressive overyielding^26^ (Extended Data Table 5). Additive partitioning revealed that the increases of net biodiversity effects were primarily driven by increases in complementarity effects (Extended Data Table 6, F_1,31_ = 9.61, P = 0.004) and weakening negative selection effects (Extended Data Table 6, F_1,37_ = 4.61, P = 0.038). In the last year of measurements, selection effects were no longer significantly different from zero (Fig. 2, F_1,31_ = 3.40, P = 0.075).

**Figure 2.**
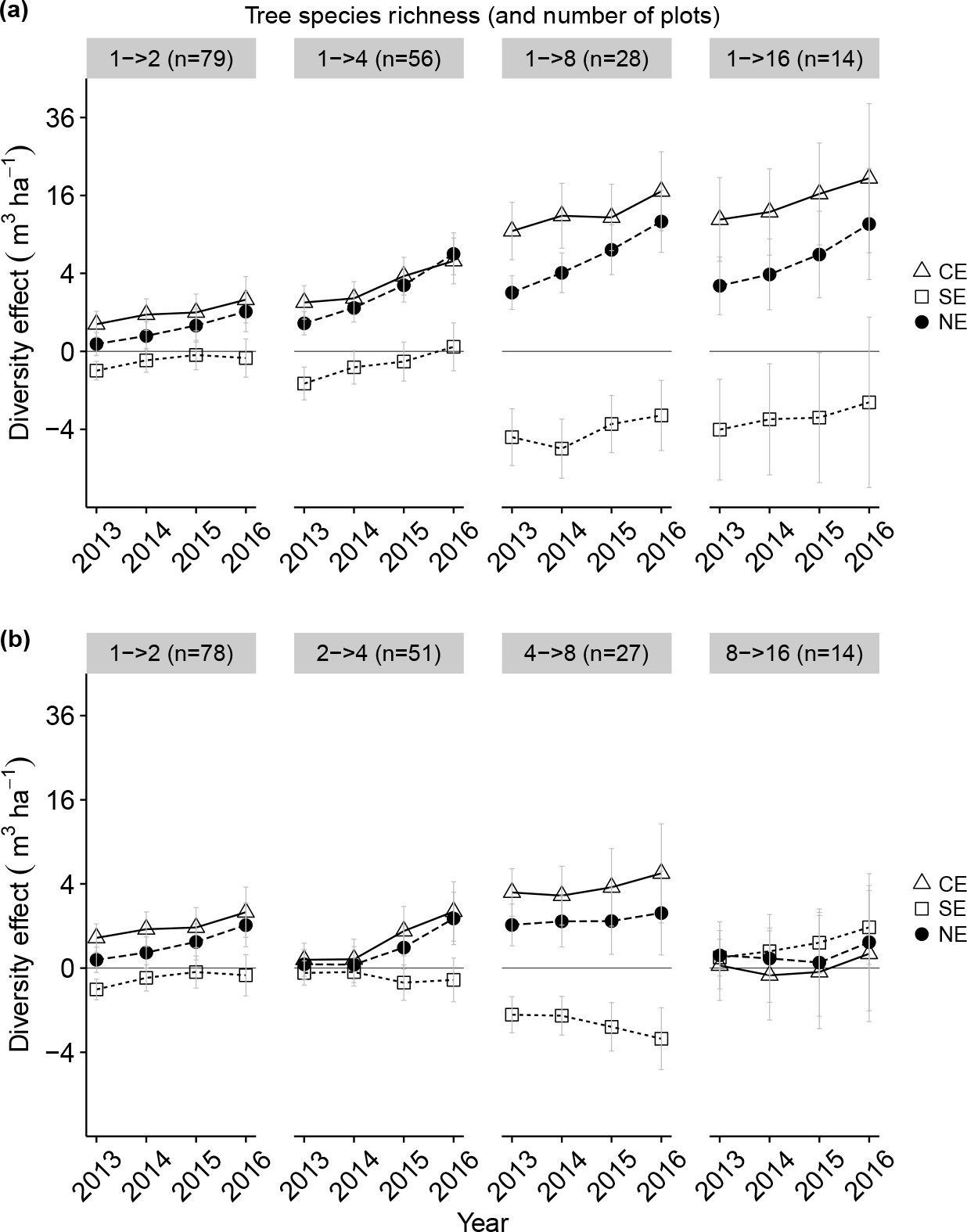
Changes over time in the net biodiversity effect (NE) and its additive components, complementarity effect (CE) and selection effect (SE), on stand-level tree volume. The figure shows means and standard errors. In (a), diversity effects were calculated with monocultures as reference (Extended Data Table 6), in (b) with component mixtures of half the number of species as reference. The y-axes are square root-scaled to reflect the quadratic nature of biodiversity effects.

We observed considerable variation in overyielding among communities of the same species-richness level. Some of this variation was explained by functional diversity but phylogenetic diversity had low explanatory power. For the 48 different 2-species mixtures, complementarity effects were positively correlated with the functional distance and selection effects with vertical crown dissimilarity, also referred to as crown complementarity between species (Fig. 3, Extended Data Table 7). That vertical crown complementarity^22^ contributed to overyielding via selection rather than complementarity effects indicated that it was due to asymmetric light competition^27^ and is consistent with the “competition-trait hierarchy hypothesis”^28^.

**Figure 3.**
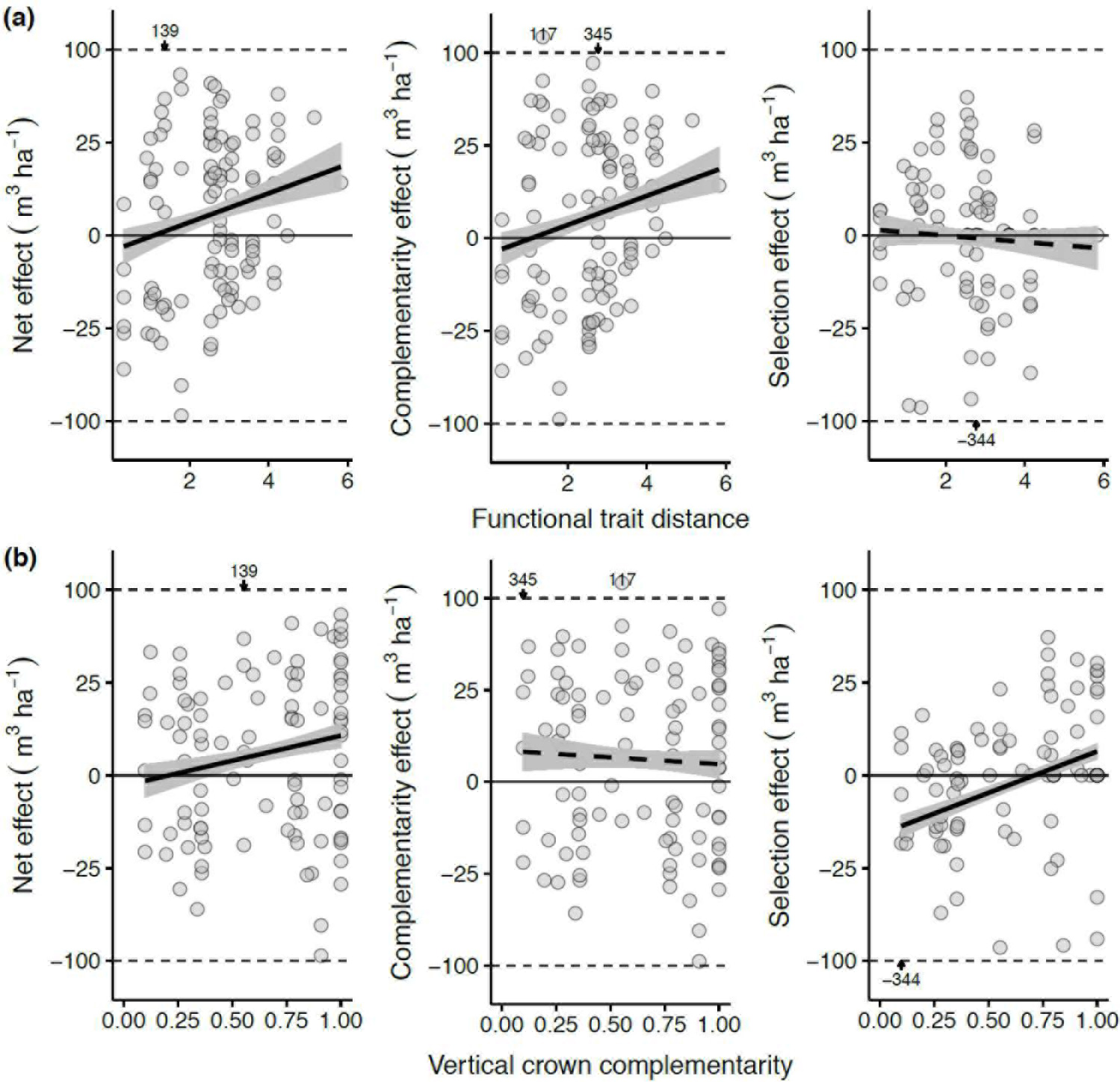
Relationships between biodiversity effects and (a) functional trait distance and (b) vertical crown complementarity (proportional dissimilarity of monoculture vertical crown extent) in 2016 (n = 108) Regression lines and confidence bands (indicating ± standard error of predicted values) are based on mixed models (Extended Data Table 7). The y-axes are square root-scaled to reflect the quadratic nature of biodiversity effects. Four extreme y-values are moved to the plot margin and given as numbers.

Species with high monoculture productivity (Fig. 4a) explained large amounts of variation in stand-level productivity (Fig. 4b), but their contribution was not always positive, as demonstrated by several negative species-level selection effects (Fig. 4c). Despite the positive effect of species richness on community productivity, the population-level responses of each species to species richness varied from positive to neutral to negative (Fig. 4d). These responses did not differ between evergreen and deciduous species (Fig. 4d, F_1,159_ = 0.89, P = 0.347). A similar decoupling between community- and population-level responses has previously been reported from grassland biodiversity experiments^8^ and indicates that a few positive population-level responses can overcompensate a larger number of negative population-level responses. Nevertheless, the number of species with positive responses to community diversity and the magnitude of their responses increased with time (Fig. 4d).

**Figure 4.**
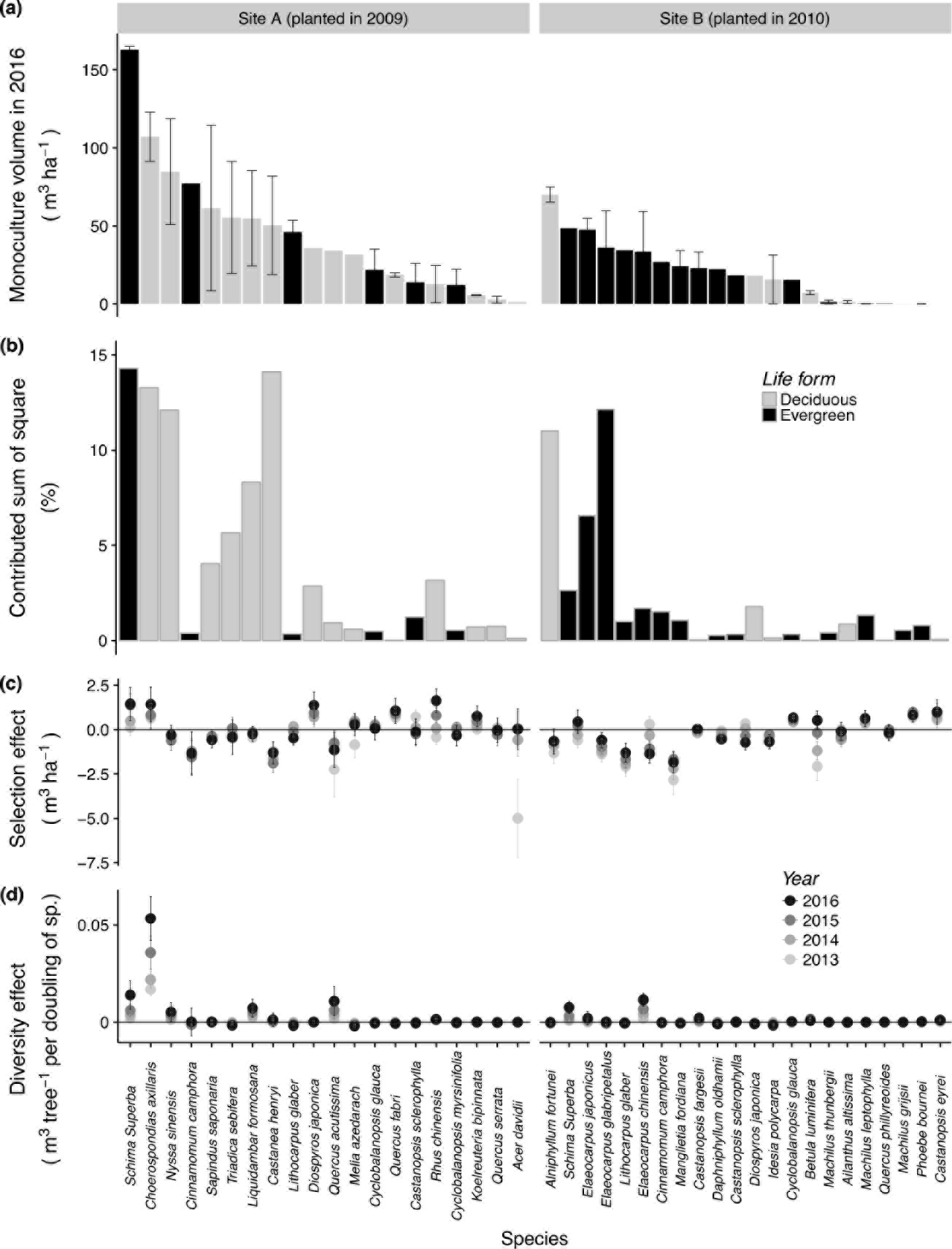
Monoculture stand-level tree volume of species in 2016 (a) and the fraction of stand-level tree volume sum of squares explained by the presence of each species in a plot (b), their species-specific selection effects (SEs) on stand-level tree volume (c) and their tree-level volume response to species richness (d). Bars indicate standard errors. For (d) the volume of each species, standardized for the number of originally planted individuals of that particular species, was linearly regressed against log_2_(tree species richness) with the data from (sub)plots without shrub species.

Competition by understorey shrubs planted in the gaps between the trees reduced stand-level tree volume (Extended Data Table 8, F_1,234_ = 4.80, P = 0.029), but this effect decreased with shrub species richness (Extended Data Table 8, F_1,499_ = 5.40, P = 0.022) and was negligible when mixtures of 8 shrub species were planted (Extended Data Fig. 6), indicating reduced competition between shrubs and trees at higher shrub diversity levels. The diversity-productivity relationships we found were scale-independent, i.e. they did not differ between 1- and 4-mu plots (Extended Data Table 8, F_1,114_ = 0.20, P = 0.694 for interaction species richness × plot size).

Our results provide strong evidence for a positive effect of tree species richness on tree productivity at stand-level in establishing subtropical forest ecosystems, and support the idea that highly diverse subtropical forest ecosystems are niche-structured^22,27^. Seven-year old mixed-species stands can produce an estimated additional aboveground wood volume of 25 m^3^ ha^−1^ relative to the average monoculture, which translates to the sequestration of approximately an extra 10 Mg C ha^−1^ (Fig. 1, Extended Data Fig.4). We expect this effect to grow further, given that we did not observed any signs of a deceleration over the present measurement period.

The size of the biodiversity effects we found for these forests is similar to biodiversity effects reported from grassland studies^8,9^. Given that plant biomass is higher in forests, and that the largest fraction of tree carbon is bound in relatively persistent woody biomass, these effects translate into significant diversity-mediated rates of carbon sequestration. Substantial forest areas are managed world-wide, with large afforestation programs underway in many countries. In China, huge economic efforts are made for afforestation, with a net growth of total forested area by 1.5×10^6^ ha yr^−1^ achieved from 2010 to 2015 ^29^. However, the overwhelming fraction of newly established forests are monoculture plantations of species with highest productivity in the short term^30^. Our analysis suggests that a similar productivity could be achieved with mixed plantations of native species, which would result in co-benefits in the form of biodiversity management and a likely higher level and stability of ecosystem services in the longer term.

## Online content

Methods, along with additional Extended Data display items and Source Data, are available in the online version of the paper; references unique to these sections appear only in the online paper.

**Extended Data** are available in the online version of the paper.

## Acknowledgements

We thank the farmers and Chen Lin for help in the field. This study was supported by the German Research Foundation (DFG FOR 891), the National Natural Science Foundation of China (NSFC No. 31270496 and No. 31300353), the Swiss National Science Foundation (SNSF No. 130720, 147092) and the European Union (EC 7th Framework Program No. 608422).

### Author contributions

HB, KM and BS conceived the project with help from all co-authors; YH carried out the measurements; YH, YC, KM, PAN and BS led the data analysis and interpretation. All authors contributed to the writing of the manuscript.

### Author information

The authors declare no competing financial interests. Correspondence and requests for materials should be addressed to Y.H., H.B., K.M., P.A.N. or B.S.

## METHODS

### Study site and experimental design

The BEF-China experimental platform was established in Jiangxi Province, China (29°08′-29°11′N, 117°90′-117°93′E). Climate at the site is subtropical, with mean annual temperature and precipitation of 16.7°C and 1800 mm, respectively (averaged from 1971-2000)^31^. A large-scale tree biodiversity experiment was established in 2009-2010 on two sites (A and B) of approximately 20 ha each, with a total of 226’400 individual trees planted. Here, we use all plots in which random species-loss scenarios were simulated. The species pool contains 40 tree species (Extended Data Table 1), 24 for each site (of which eight are shared between sites). The 24 species at each site were divided into three 8-species sets. By combing these 8-species sets in all possible ways, three pools of 16 species were created. The species in each 16-species pool were put in random sequence and then repeatedly divided in halves until monocultures were obtained. This procedure resulted in 70 unique species compositions per site (Extended Data Table 2) and ensured that each tree species occurred in equal overall proportion at each diversity level. We further included monoculture plots with two commercially important tree species, *Pinus massoniana* and *Cunninghamia lanceolata*, as reference, with 5 replicate plots per species and site. Each plot was 25.8 × 25.8 m in size and planted with 400 tree individuals arranged on a rectangular 20 × 20 grid with 1.29 m spacing between rows and columns. To minimize edge effects, plots were established adjacent to each other, with trees thus forming a continuous cover across the entire site. Site A was planted in 2009, site B in 2010.

Plots of one species pool per site (pools A1 and B1 at sites A and B, respectively, Extended Table 2) were additionally replicated in plots that were four times larger and thus contained 1600 trees. These large plots were subdivided into four quadrants in which a factorial understorey shrub-diversity treatment was established. These four subplots either had no shrub understorey (0 species), or shrubs planted in all the centers between 4 adjacent trees, at a diversity of 2, 4 or 8 shrub species (Fig. 1a).

The design we use here consisted of 140 small plots (1 mu) and 64 large plots (4 mu). Out of this total of 396 1-mu sized (sub)plots, nine had to be excluded because these were not established due to a lack of sapling material or high initial mortality. All plots were weeded annually to remove emerging herbs and woody species that were not part of the planting design.

### Tree measurements

We assessed stand-level and population-level tree growth by measuring the height of trees and maximum and minimum stem diameter at 5 cm above ground to calculate basal area. We focused on the central 4 × 4 =16 trees of each 1-mu (sub)plot to avoid edge effects. These measurements were repeated annually in September/October from 2013 to 2016. We aggregated these tree-level data at the species (i.e. population) and stand level.

We further calculated a cylindrical tree volume as the product of basal area and height. The true volume was then obtained by multiplying this proxy with a form factor determined by a complete harvest of 154 trees in natural forest near the experimental sites. The total volume of each harvested trees was calculated as ratio of total aboveground dry biomass and average wood density. Similarly, tree biomass was determined by multiplying the cylindrical volume of each experimental tree with a biomass conversion factor determined based on the harvested trees (Extended Data). Biomass was converted to carbon content^32^ by multiplying with 0.474 g C g^−1^.

### Complementarity effect and selection effect

We used the additive partitioning method of Loreau & Hector^3^ to decompose net biodiversity effects (NEs) of productivity measures into complementarity (CEs) and selection effects (SEs), separately for each year and diversity level. CEs and SEs depend on relative yields of species, which we calculated using monoculture biomass as denominator. If a species failed to establish in monoculture (which was the case for *Meliosma flexuosa*, *Castanopsis eyrei* and *Machilus grijsii*), or had a mortality exceeding 80% (*Quercus phillyreoides*, *Phoebe bournei*), it was excluded from the set of target species in the corresponding mixtures^33^. Formally, CEs and SEs are related to (co)variances and therefore were square-root transformed with sign reconstruction 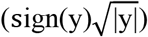 prior to analysis, which improved the normality of residuals^3^.

### Overyielding and underyielding

Overyielding describes the case where the productivity of a mixture exceeds the average productivity of monocultures of component trees^26^. Conversely, underyielding identifies a lower yield of the mixture relative to monocultures. Transgressive overyielding indicates that the productivity of a mixture exceeds the productivity of the monoculture of the most productive component species. Transgressive underyielding is defined similarly. We determined overyielding and underyielding of all mixtures relative to monocultures. Capitalizing on the nested nature of our design, we further determined the same metrics using the two mixtures with half the set of species as reference, instead of monocultures, i.e. we tested whether combining communities with two sets of species resulted in a community that produced more or less biomass than expected on the assumption of no interactions among the sets (overyielding) or that community productivity would be determined by the more productive set of species alone (transgressive overyielding).

### Vertical crown complementarity

We quantified the interspecific complementarity in vertical crown extent of trees in 2016. The crown extent was determined as interval between the lowest side-branch and the top of a tree in monocultures. These data were averaged across all surviving trees of the 16 central individuals planted in a plot. We then calculated vertical crown complementarity in 2-species mixtures as proportional dissimilarity of the crown extents between the two species:

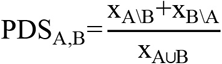

where x_A\B_ indicates the vertical extent (in meters) that is occupied by A but not by B (vice versa for x_B\A_), and x_A⋃B_ indicates the extent occupied by at least one of the species. This index is equivalent to one minus the proportional similarity index proposed by Colwell and Futuyma^34^.

### Statistical analysis

We used analysis of variance based on type-I sum of squares linear mixed-effects models to assess the effects of tree species richness (and additional design variables) on productivity^35^. All analyses were done in R 3.3.2 and ASReml-R^36^. The models included the fixed effects site, tree species richness (log_2_-transformed), year (continuous variable, centered over our observation period), the interaction log_2_(tree species richness) × year, and the interaction site × year. Random effects were species composition (with a separate variance component for each site), plot (with a separate variance component for each site), subplot, and the interactions of all these random terms with year. Model residuals were checked for normality and homogeneity of variances.

For the analyses of shrub diversity effects, the model contained the additional fixed effects shrub presence (a two-level factor: 0 vs. 2, 4 or 8 shrub species), plot size (a two-level factor: 1 vs. 4 mu), log2 of shrub species richness (for shrub-species richness >0), and the interactions of all these terms with log2(tree species richness) and with year. Random effects were species composition (with a separate variance component for each site), plot (with a separate variance component for each site), subplot, and the interactions of all these random terms with year (Extended Data Table 6). The interaction of year and site and the site-specific variance terms estimated for some random terms accounted for the fact that site B was established one year after site A and that trees at site B were therefore smaller.

## Data availability statement

The data supporting the findings of this study will be deposited in Pangaea with the accession code https://doi.pangaea.de/xxxxxxxxxxxxxx.

## EXTENDED DATA

### Conversion factors for volume, biomass and carbon content

We harvested 154 trees in a natural forest in 2010 near the experimental sites to determine conversion factors from cylindrical volume (tree basal area × height) to true volume and biomass. The trees belonged to eight common species and three life forms (evergreen, deciduous and coniferous) and were chosen to represent a naturally occurring size span of young trees.

Trees were separated into large woody parts (stems and large branches with a diameter ≥ 3 cm), twigs (the apical part of the stem and large branches plus side branches with a diameter < 3 cm), and dead attached material (large dead branches or twigs). Branches were divided into segments of typically about 1 m length. The volume of large woody parts and twigs was determined geometrically, approximating the parts as truncated cone (large woody parts, 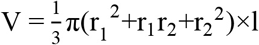 where l is the length and r_1_ and r_2_ are the end radius), or cone (twigs, as above but r_2_=0). The density of these fractions was determined by oven-drying a representative subsample of stem and branch discs or twigs.

These geometric and density data were then scaled up to total aboveground tree biomass using a Bayesian framework, modeling twig mass and density in dependence of branch positions within tree crowns^37^.

Conversion factors from cylindrical volume to true volume (and mass) were determined as total tree volume (and tree mass, including leaves) divided by cylindrical volume. We analyzed the variation of these conversion factors with tree size and species life form using mixed effects models with species identity as random term. We found that large trees deviated from the linear relationship of form factor and cylindrical volume, and we therefore removed trees with a cylindrical volume ≥ 500 liter from the form factor calibration, leaving a set of 119 trees. Within this set, there was only a small variance among species and no significant effect of life form on the form factor; the form factor decreased linearly with the cylindrical volume of harvested trees (Extended Data Table 3). We therefore used a form factor of 0.5412 m^3^ m^−3^ − 0.1985 m^−3^.BA.h (with basal area BA in m^2^ and height h in m). The intercept of 0.5412 m^3^ m^−3^ is the weighted average form factor of evergreen and deciduous species at size zero (in our study, 19 of 40 species were evergreen and 21 deciduous). Biomass factors were determined similarly, yielding a conversion factor of 269.13 kg m^−3^ −141.96 kg m^−3^.BA.h. For the two coniferous species that were planted for comparison in monocultures only, we used separate equations obtained from the harvested trees of the same species *Pinus massoniana* and *Cunninghamia lanceolata*. Here the form factor was 0.5083 m^3^ m^−3^ − 0.1985 m^−3^.BA.h and the biomass factor was 216.79 kg m^−3^ −141.96 kg m^−3^.BA.h.

### Functional trait and phylogenetic distances

We used four functional traits related to the resource-use strategies of tree species: specific leaf area^38^, branch-wood density^38^, relative volume growth rate (RGR) and life form (deciduous or evergreen). These traits were determined in plots that were part of the experiment. RGR was calculated as the log-transformed relative difference in stand volume of monocultures between seven (2015 for site A and 2016 for site B) and five years (2013 for site A and 2014 for site B) after planting. We selected the monocultures without shrub treatments. We used site-specific RGR because of the large variation in growth rates between sites A and B. We calculated functional trait distances among species pairs in 2-species communities as Euclidean distances in standardized multivariate trait space (using the four traits as axes).

We calculated phylogenetic distances among species pairs as their cophenetic distance in a node age-calibrated phylogenetic tree^39^.

We assessed the effects of trait and phylogenetic distances on different components of diversity effects of two-species mixtures with linear mixed-effects models, where we set site and trait/phylogenetic distance as fixed effects, community composition and plot as random effect (with a separate variance component for each site). Measures of diversity effects were square-root transformed with sign reconstruction to improve normality of model residuals.

**Extended Data Figure 1.**
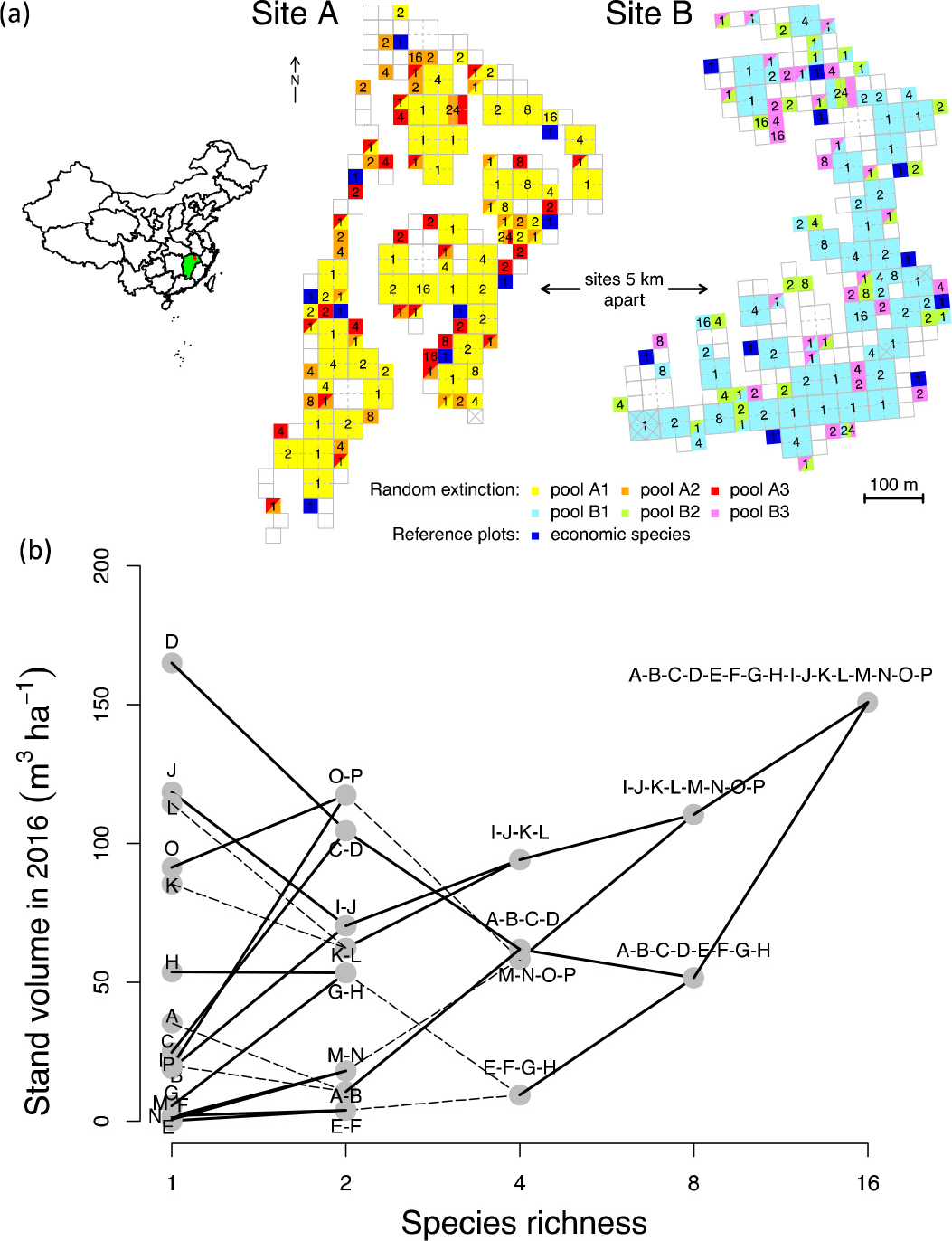
Map of BEF-China position and experimental plots of random extinction scenarios and economic trees (a). Results from species pool A1 to illustrate the “broken stick” design (b). Letters represent different species (A= *Cyclobalanopsis glauca*; B = *Quercus fabri*; C = *Rhus chinensis*; D = *Schima superba*; E = *Castanopsis eyrei*; F = *Cyclobalanopsis myrsinifolia*; G = *Koelreuteria bipinnata*; H = *Lithocarpus glaber*; I = *Castanea henryi*; J = *Nyssa sinensis*; K = *Liquidambar formosana*; L = *Sapindus saponaria*; M = *Castanopsis sclerophylla*; N = *Quercus serrata*; O = *Choerospondias axillaris*; P = *Triadica sebifera*). Solid lines represent overyielding, while dashed lines represent underyielding.

**Extended Data Figure 2.**
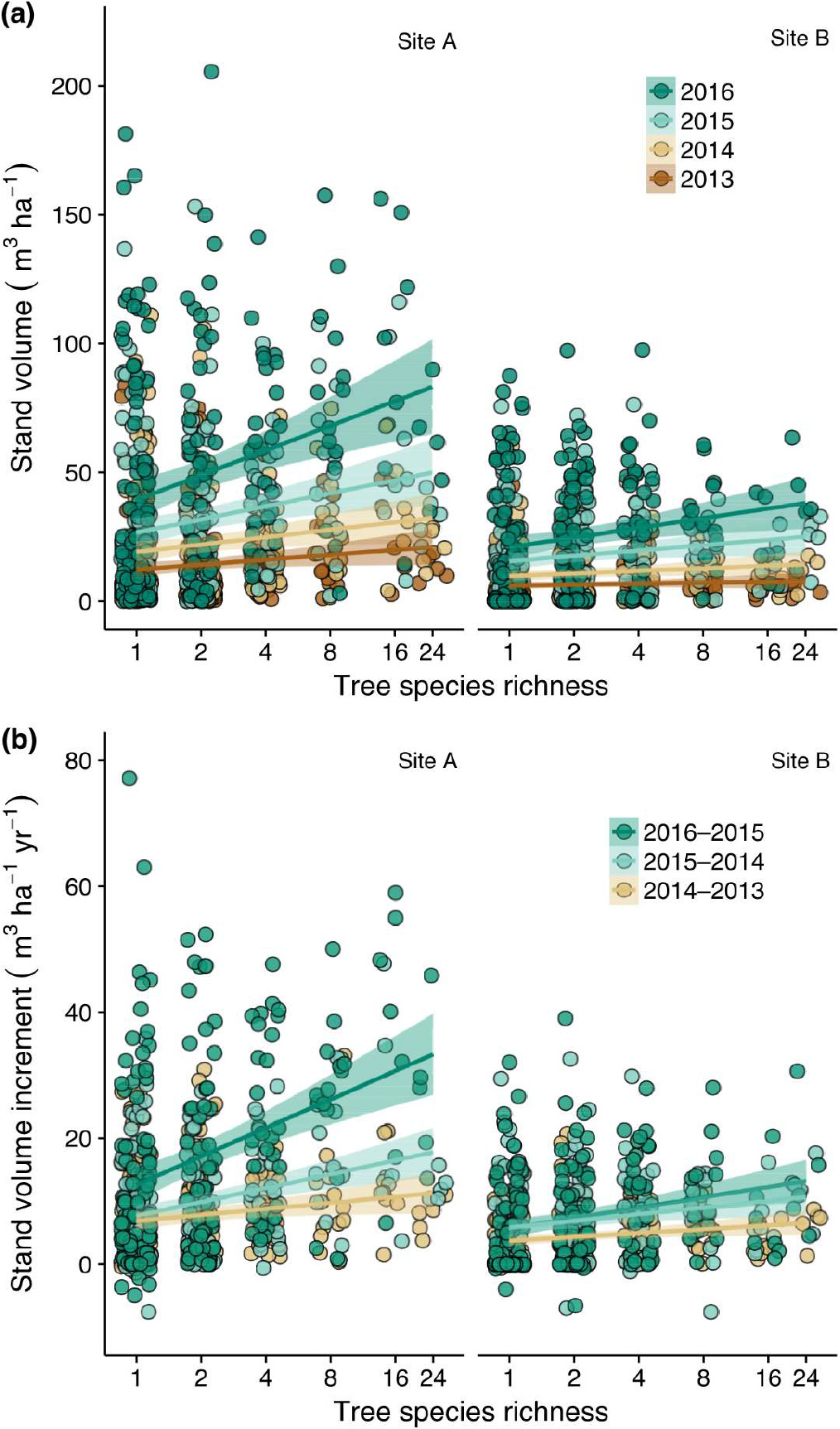
Stand-level tree volume (a) and its increment (b) as a function of tree species richness from 2013–2016. Positive effects of tree species richness increase with time. Raw data points, regression lines and 95% confidence bands are shown for each year. Note that the extremes of the point cloud necessarily taper off towards higher diversity levels for statistical rather than biological reasons; this is due to the fact that for a given diversity level extreme values are more extreme the larger the sample size is^26^.

**Extended Data Figure 3.**
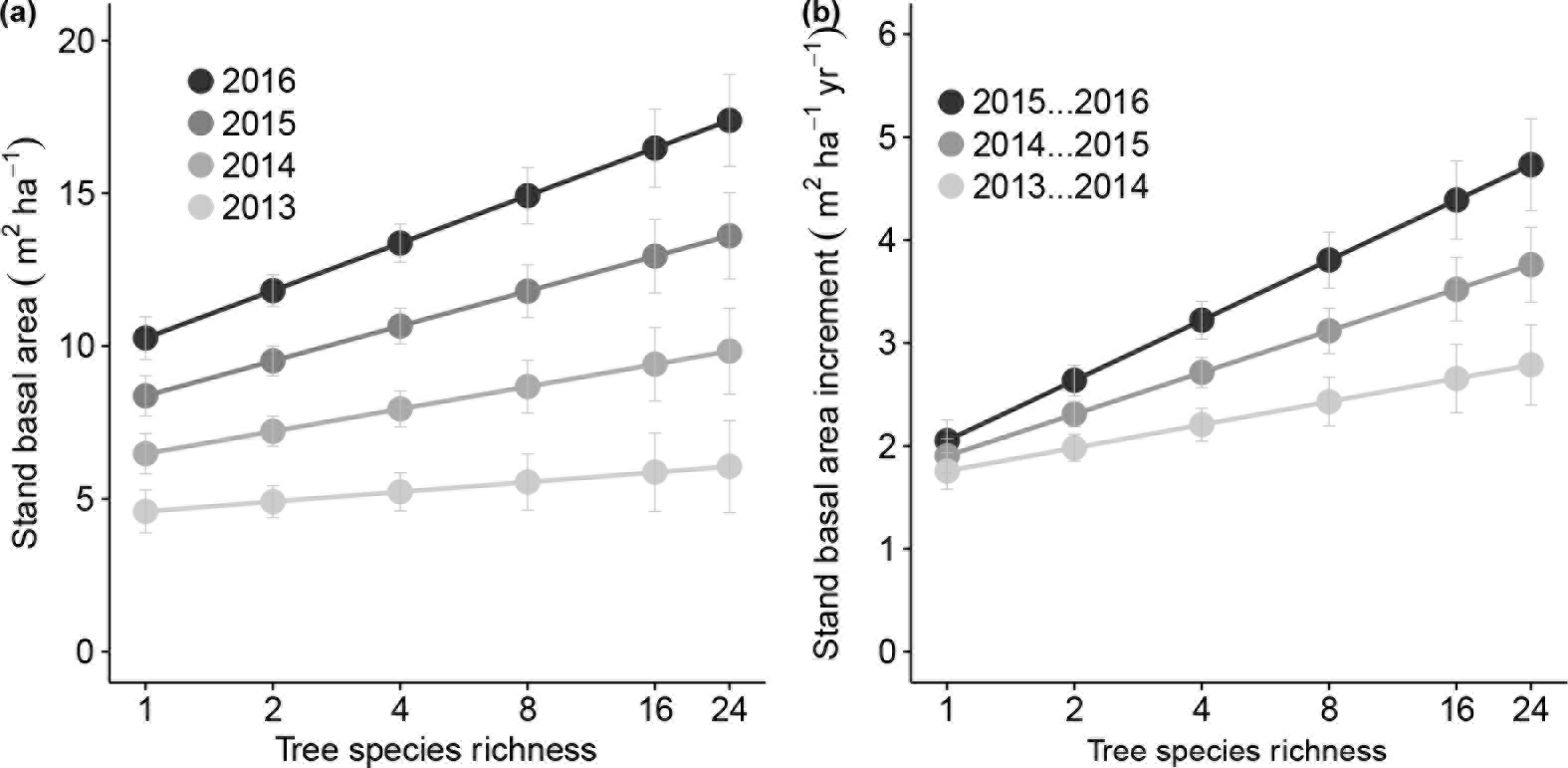
Stand-level tree basal area (a) and its annual increment (b) as a function of tree species richness from 2013–2016. The figure shows predicted means and standard errors based on fitted mixed models (Table 1). Effects of species richness were significantly positive and increased throughout the observation period.

**Extended Data Figure 4.**
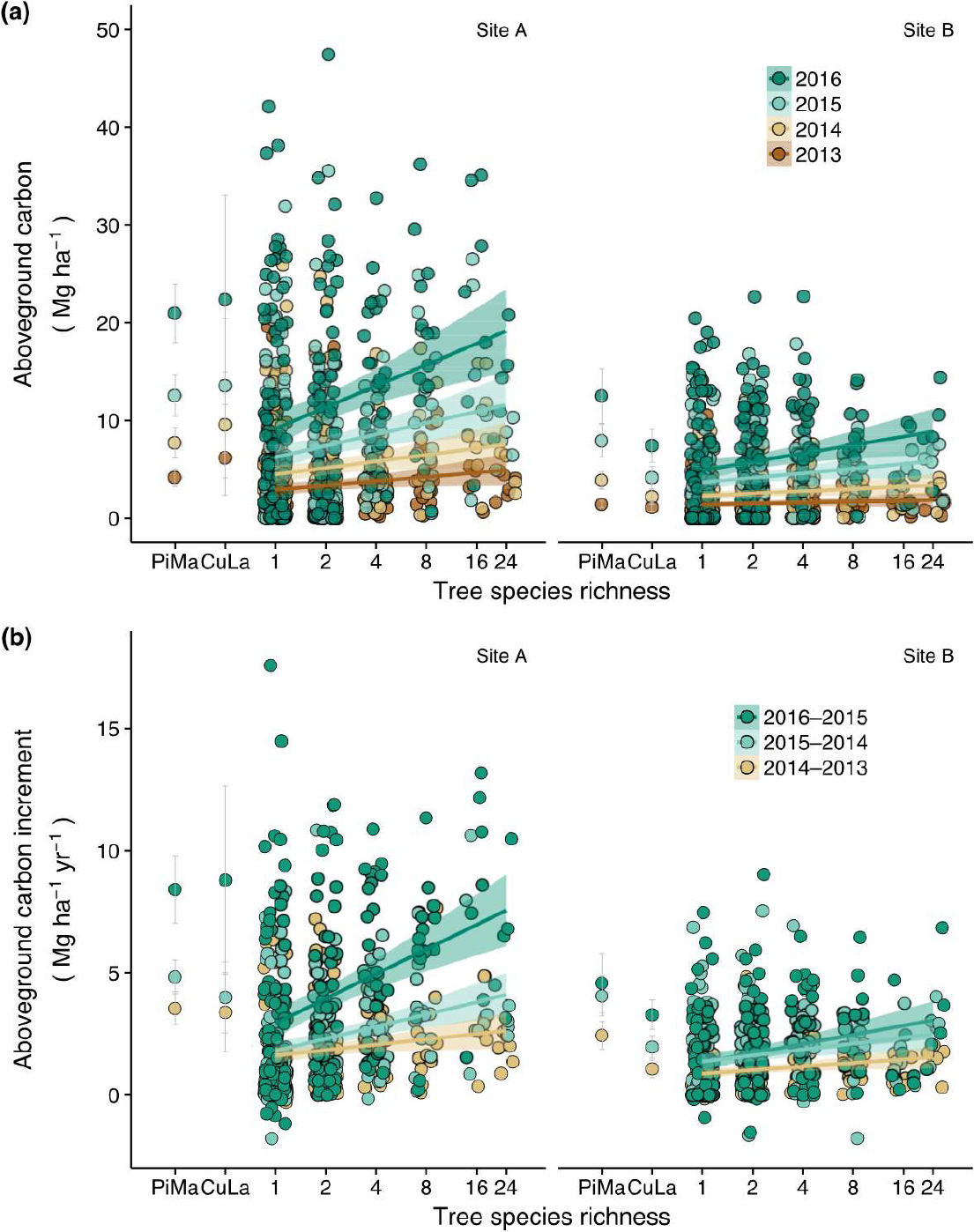
Aboveground stand-level tree carbon (a) and its annual increment (b) as a function of tree species richness from 2013–2016. Raw data points, regression lines and 95% confidence bands are shown. On the left of each panel means ± standard errors for the two economic tree species (PiMa = *Pinus massoniana*; CuLa = *Cunninghamia lanceolata*) are inserted. Note that the extremes of the point cloud necessarily taper off towards higher diversity levels for statistical reasons; this is due to the fact that for a given diversity level extreme values are more extreme the larger the sample size is^26^.

**Extended Data Figure 5.**
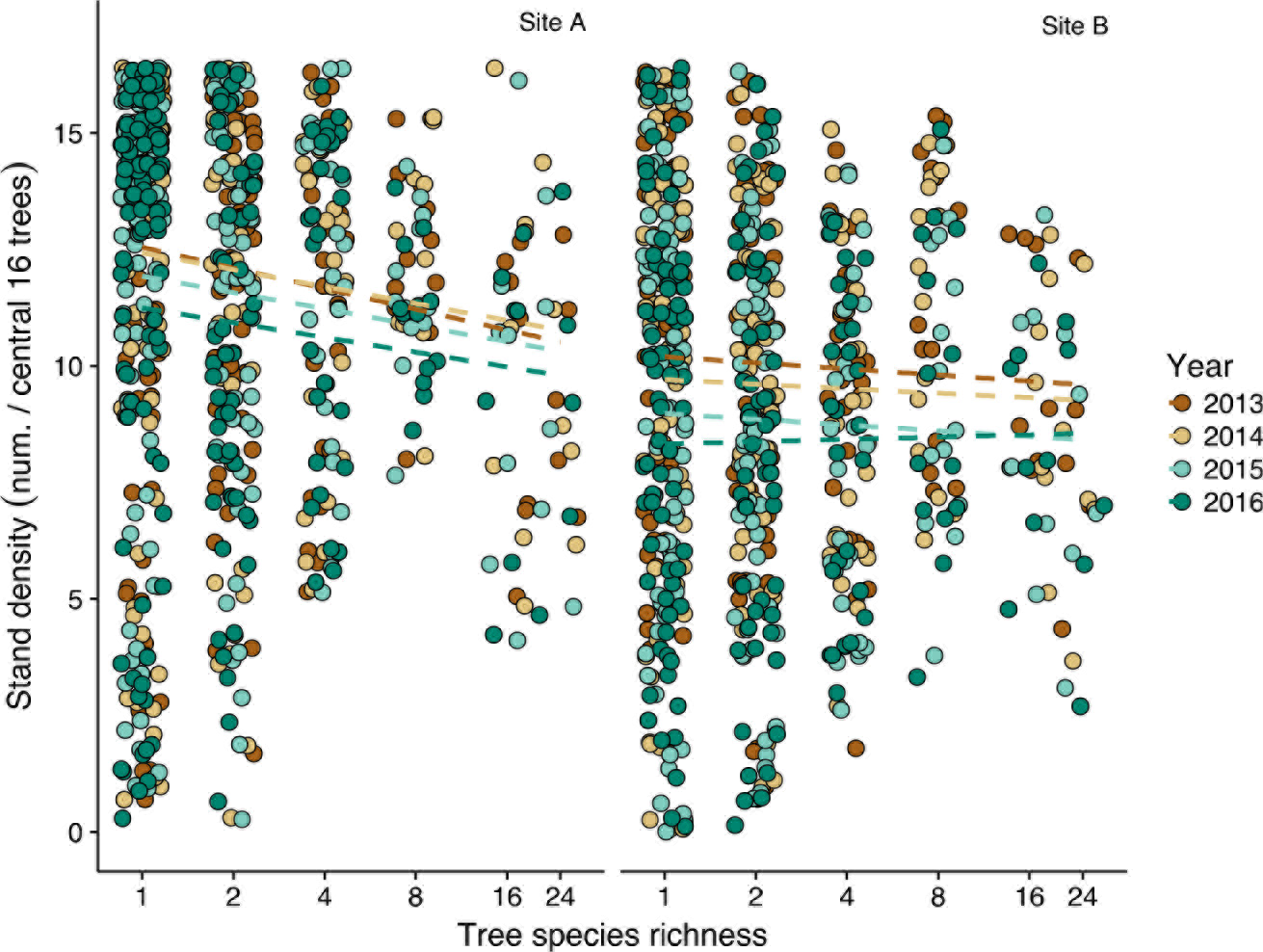
Stand density as a function of tree species richness from 2013–2016. Raw data points together with non-significant regression lines (dashed) are shown. Density indicates the number of surviving trees out of 16 planted in the central area of each plot.

**Extended Data Figure 6.**
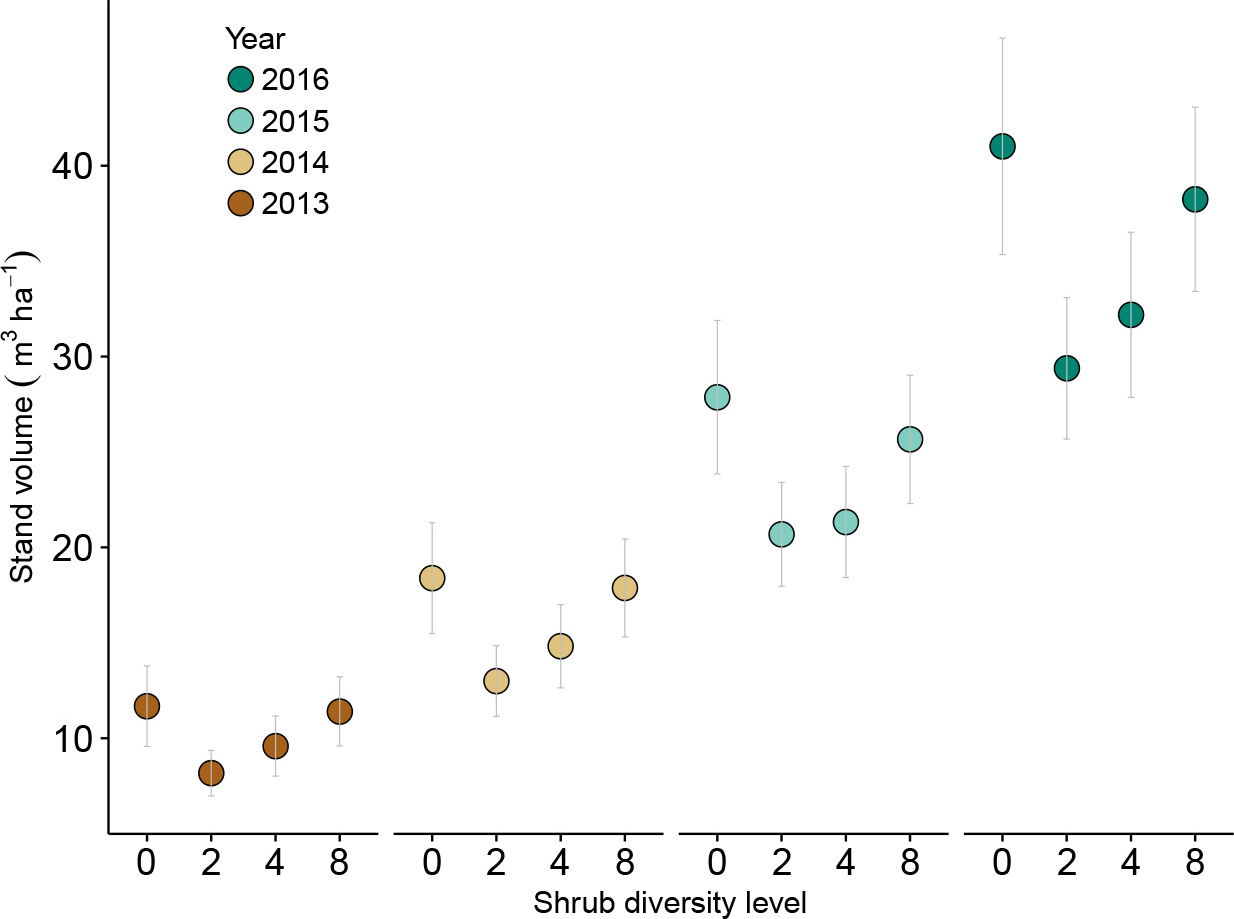
Effects of shrub diversity on average stand-level tree volume in species pools A1 and B1. Data are from 4mu plots. The figure shows predicted means and standard errors based on a fitted mixed model (Extended Data Table 8).

**Extended Data Table 1.**
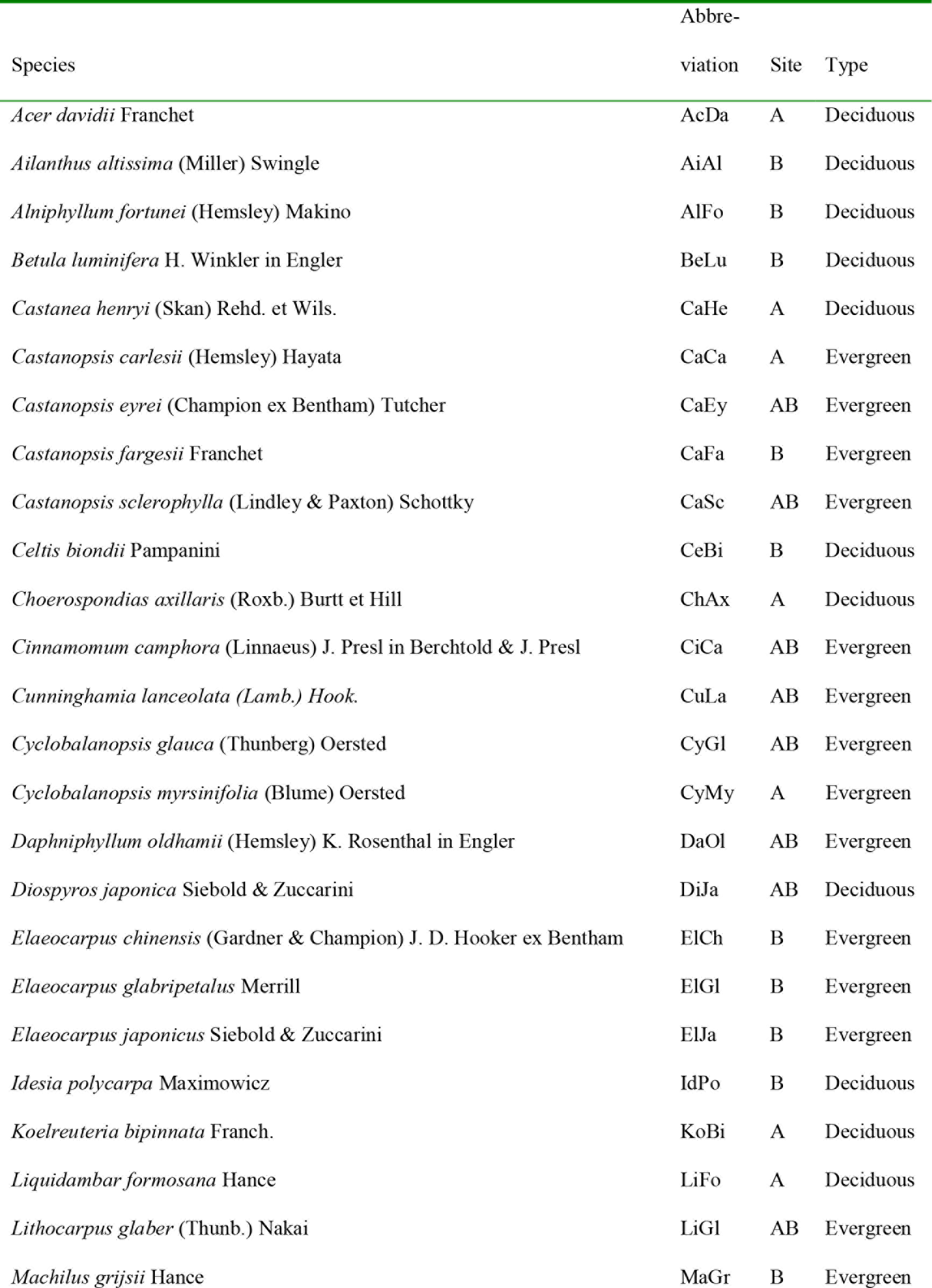
List of tree species used in the BEF-China experiment according to the Flora of China (http://www.efloras.org and http://frps.eflora.cn).

Notes:

The site column shows the experimental site (A, B) where the species was planted. The type column shows species life form (D = deciduous species; E = evergreen species).

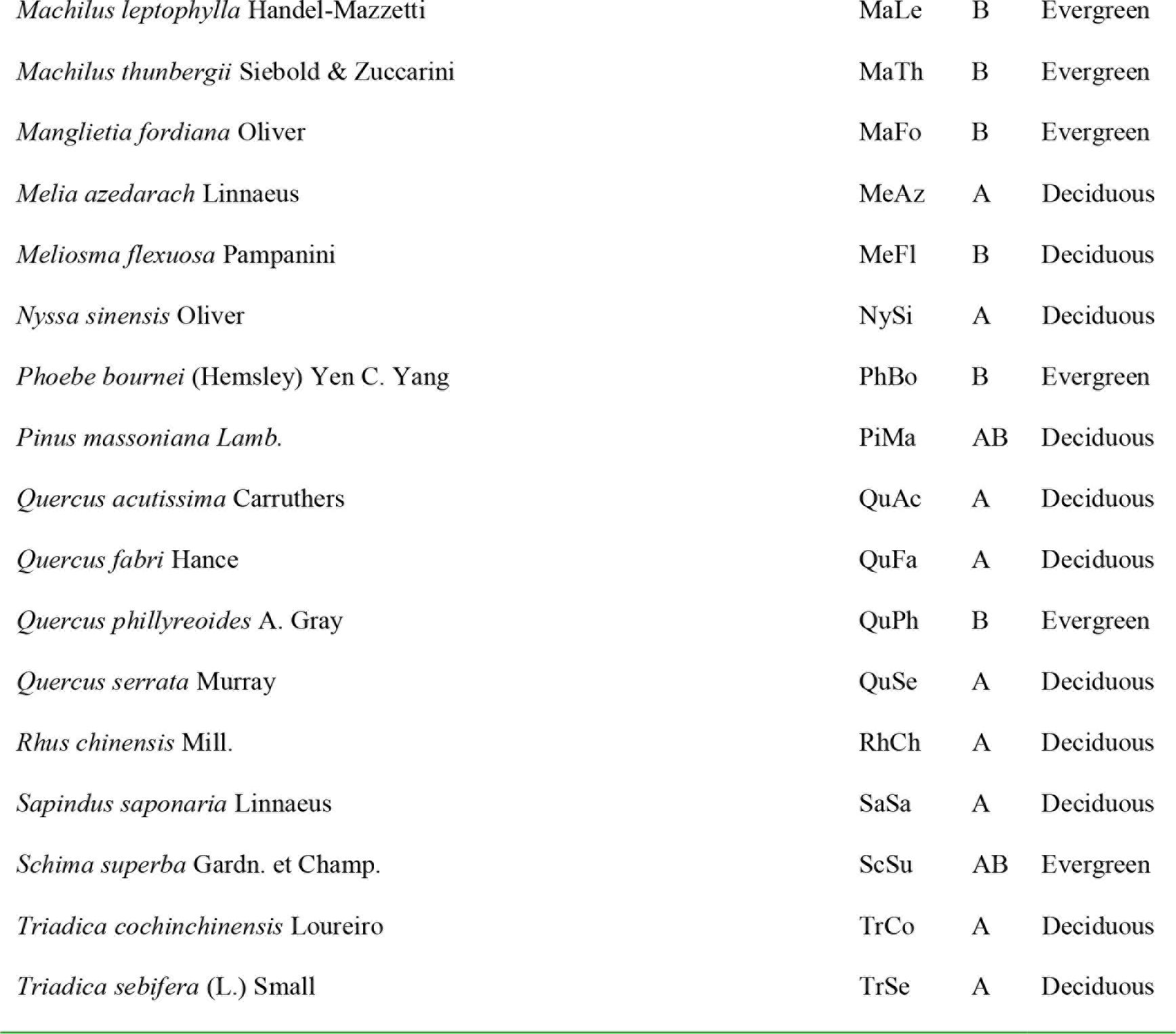

**Extended Data Table 2.**
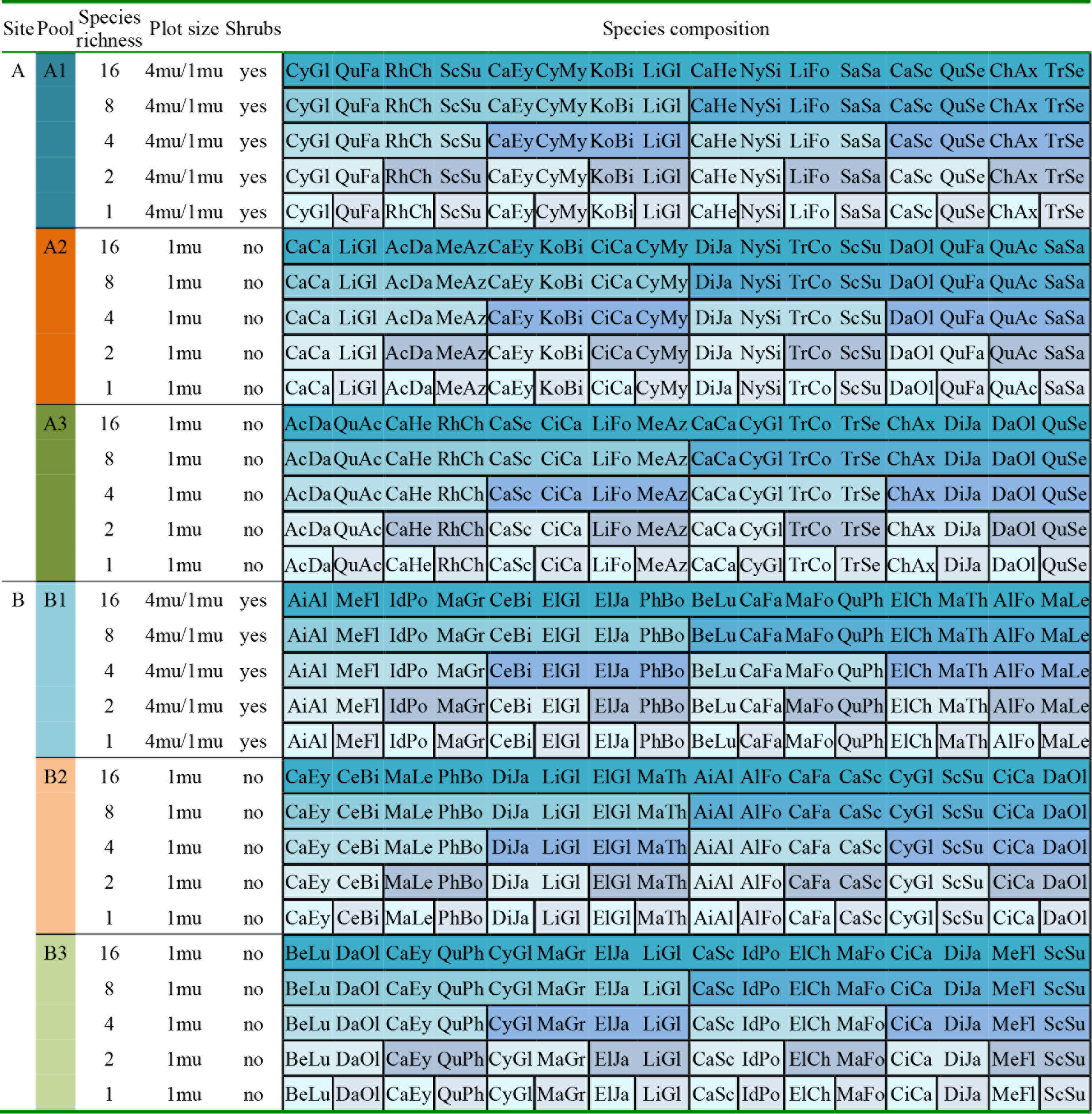
Experimental design.

Note:

See Extended Data Table 1 for species abbreviations.

**Extended Data Table 3.** Mixed-effects model for the effects of cylindrical volume and life form on form and biomass factors.

|  | Form factor |  |  |  | Biomass factor |  |  |  |
| --- | --- | --- | --- | --- | --- | --- | --- | --- |
|  | Df | ddf | *F* | *P* | ddf | denDF | *F* | *P* |
| (Intercept) | 1 | 4.4 | 2418 | <0.001 | 1 | 4.8 | 828.9 | <0.001 |
| Cylindrical volume | 1 | 105.7 | 8 | 0.007 | 1 | 114.7 | 11.2 | 0.001 |
| Life form | 2 | 4.7 | 20 | 0.005 | 2 | 4.9 | 22.1 | 0.004 |
| Variance components | Component |  | s.e. | z-ratio | Component |  | s.e. | z-ratio |
| Species identity | 0.00002 |  | 0.00058 | 0.043 | 257.90 |  | 356.87 | 0.723 |
| Residual | 0.01237 |  | 0.00169 | 7.321 | 4384.52 |  | 590.66 | 7.423 |

Notes:

Notes:

Fixed effects were fitted sequentially (type-I sum of squares) as indicated in the table (the random term was species identity). Abbreviations: df= nominator degree of freedom; ddf = denominator degree of freedom; s.e. = standard error; *F* and *P* indicate F-ratios and P-values of the significance tests.

**Extended Data Table 4.** Mixed-effects models for the effects of site, tree species richness (logSR), time (year) and interactions on aboveground stand-level tree carbon and its increment.

| Source of variation | Aboveground carbon |  |  |  | Aboveground carbon increment |  |  |  |
| --- | --- | --- | --- | --- | --- | --- | --- | --- |
|  | df | ddf | *F* | *P* | df | ddf | *F* | *P* |
| Site | 1 | 101.30 | 19.12 | <0.001 | 1 | 104.70 | 14.42 | <0.001 |
| LogSR | 1 | 89.20 | 5.16 | 0.026 | 1 | 94.60 | 9.11 | 0.003 |
| Year | 1 | 104.20 | 209.50 | <0.001 | 1 | 106.20 | 70.30 | <0.001 |
| Site × year | 1 | 104.80 | 18.47 | <0.001 | 1 | 110.60 | 26.78 | <0.001 |
| LogSR × year | 1 | 95.90 | 10.54 | 0.002 | 1 | 97.00 | 11.63 | <0.001 |

Notes:

Notes:

Fixed effects were fitted sequentially (type I sum of squares) as indicated in the table (random terms were community composition, plot, subplot and the interaction of these with year, with site-specific variance components for species composition and plot). Abbreviations: df= nominator degree of freedom; ddf = denominator degree of freedom; logSR = log_2_(tree species richness). *F* and *P* indicate F-ratios and P-values of the significance tests.

**Extended Data Table 5a.** Average number of 1-mu (sub)plots with overyielding (Over) and underyielding (Under) for stand-level tree volume in 2016 across richness levels.

| Reference | Target -><br>reference | All plots |  |  | Transgressive plots |  |  |
| --- | --- | --- | --- | --- | --- | --- | --- |
|  |  | Over | Under | *P*<br>(over>under) | Over | Under | *P*<br>(over>under) |
| Component<br>monocultures | 2->1 | 65 | 47 | 0.088 | 43 | 21 | 0.005 |
|  | 4->1 | 39 | 17 | 0.003 | 14 | 0 | <0.001 |
|  | 8->1 | 21 | 7 | 0.007 | 3 | 0 | 0.041 |
|  | 16->1 | 9 | 5 | 0.280 | 1 | 0 | 0.239 |
|  | total | 134 | 76 | <0.001 | 62 | 21 | <0.001 |
| Component | 2->1 | 65 | 47 | 0.088 | 43 | 22 | 0.009 |
| mixtures with | 4->2 | 32 | 24 | 0.284 | 20 | 9 | 0.039 |
| half the | 8->4 | 17 | 11 | 0.250 | 10 | 6 | 0.315 |
| number of | 16->8 | 7 | 7 | 1.000 | 6 | 5 | 0.763 |
| species | total | 121 | 89 | 0.027 | 79 | 42 | <0.001 |

**Extended Data Table 5b.** Average number of 1-mu (sub)plots with overyielding (Over) and underyielding (Under) for stand-level tree volume in different years.

| Reference | Year | All plots |  |  | Transgressive plots |  |  |
| --- | --- | --- | --- | --- | --- | --- | --- |
|  |  | Over | Under | *P*<br>(over>under) | Over | Under | *P*<br>(over>under) |
| Component monocultures | 2013 | 126 | 84 | 0.004 | 50 | 20 | <0.001 |
|  | 2014 | 128 | 82 | 0.001 | 61 | 21 | <0.001 |
|  | 2015 | 133 | 77 | <0.001 | 60 | 24 | <0.001 |
|  | 2016 | 134 | 76 | <0.001 | 61 | 21 | <0.001 |
| Component mixtures with half the number of species | 2013 | 112 | 98 | 0.33 | 69 | 37 | 0.002 |
|  | 2014 | 111 | 99 | 0.41 | 74 | 40 | 0.001 |
|  | 2015 | 115 | 95 | 0.17 | 73 | 43 | 0.005 |
|  | 2016 | 121 | 89 | 0.027 | 79 | 42 | <0.001 |

Notes:

Notes:

*P*-values indicate significance of differences between the numbers of overyielding vs. underyielding plots (χ^2^-test), or between transgressively overyielding vs. transgressively underyielding plots.

**Extended Data Table 6.** Mixed-effects models for the effects of site, tree species richness (logSR), time (year) and the interaction of the latter two on the biodiversity effects NE, CE and SE.

|  | NE |  |  |  | CE |  |  |  | SE |  |  |  |
| --- | --- | --- | --- | --- | --- | --- | --- | --- | --- | --- | --- | --- |
|  | df | ddf | *F* | *P* | df | ddf | *F* | *P* | df | ddf | *F* | *P* |
| Intercept | 1 | 60.7 | 20.43 | <0.001 | 1 | 50.7 | 43.54 | <0.001 | 1 | 52.7 | 9.29 | 0.004 |
| Site | 1 | 64.5 | 0.18 | 0.675 | 1 | 58.9 | 2.69 | 0.107 | 1 | 59.7 | 3.74 | 0.058 |
| LogSR | 1 | 60.5 | 4.58 | 0.036 | 1 | 50.6 | 9.79 | 0.003 | 1 | 52.6 | 5.89 | 0.019 |
| Year | 1 | 47.7 | 23.61 | <0.001 | 1 | 31.3 | 9.61 | 0.004 | 1 | 37.3 | 4.61 | 0.038 |
| LogSR × year | 1 | 47.1 | 0.81 | 0.374 | 1 | 31.1 | 0.26 | 0.617 | 1 | 37.0 | 0.24 | 0.627 |

Notes:

Notes:

Biodiversity effects were square-root transformed with sign reconstruction 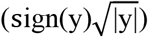. Fixed effects were fitted sequentially (type-I sum of squares) as indicated in the table (random terms were community composition, plot, subplot and the interaction of these with year, with site-specific variance components for species composition and plot). Abbreviations: df= nominator degree of freedom; ddf = denominator degree of freedom. *F* and *P* indicate F-ratios and P-values of the significance tests. The first line “Intercept” shows that the overall mean for all biodiversity effects differs significantly from zero (positively for NE and CE, negatively for SE).

**Extended Data Table 7.** Mixed-effects models for the effects of functional distance (FD), phylogenetic distance (PD) or vertical crown complementarity (PDS) on the biodiversity effects NE, CE and SE in 2-species tree stands.

|  | NE |  |  |  | CE |  |  |  | SE |  |  |  |
| --- | --- | --- | --- | --- | --- | --- | --- | --- | --- | --- | --- | --- |
|  | df | ddf | *F* | *P* | df | ddf | *F* | *P* | df | ddf | *F* | *P* |
| FD | 1 | 34.6 | 4.34 | 0.045 | 1 | 17.3 | 5.09 | 0.037 | 1 | 94 | 0.28 | 0.600 |
| PD | 1 | 58.4 | 0.92 | 0.342 | 1 | 76.3 | 0.07 | 0.787 | 1 | 39.4 | 0.67 | 0.420 |
| PDS | 1 | 22.5 | 3.43 | 0.077 | 1 | 56.5 | 0.20 | 0.659 | 1 | 8.2 | 26.43 | < 0.001 |

Notes:

Notes:

Biodiversity effects were square-root transformed with sign reconstruction 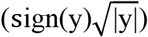. The effects of FD, PD and PDS were fitted after site (random terms were species composition and plot, considering a separate variance component for each site). Abbreviations: df= nominator degree of freedom; ddf = denominator degree of freedom. F and P indicate F-ratios and P-values of the significance tests.

**Extended Data Table 8.** Mixed-effects model for the effects of site, tree species richness (logSR), shrub presence, plot size, shrub species richness (logShrubSR), time (year) and interactions on stand-level tree volume. Data are from species pool A1 and B1, which include a shrub treatment in the planting design.

| Source of variation | df | ddf | *F* | *P* |
| --- | --- | --- | --- | --- |
| Site | 1 | 46.2 | 12.20 | 0.001 |
| LogSR | 1 | 45.0 | 3.70 | 0.059 |
| Shrub presence | 1 | 233.5 | 4.80 | 0.029 |
| Plot size | 1 | 116.2 | 0.90 | 0.353 |
| LogShrubSR | 1 | 198.8 | 5.40 | 0.022 |
| Year | 1 | 46.9 | 86.80 | <0.001 |
| LogSR × shrub presence | 1 | 232.2 | 0.80 | 0.380 |
| LogSR × plot size | 1 | 114.2 | 0.20 | 0.694 |
| LogSR × logShrubSR | 1 | 198.8 | 0.30 | 0.606 |
| LogSR × year | 1 | 46.9 | 7.20 | 0.010 |
| Shrub presence × year | 1 | 235.1 | 9.50 | 0.002 |
| Plot size × year | 1 | 128.7 | 0.00 | 0.942 |
| LogShrubSR × year | 1 | 197.1 | 3.30 | 0.069 |

Notes:

Notes:

Fixed effects were fitted sequentially (type-I sum of squares) as indicated in the table (random terms were community composition, plot, subplot and the interaction of these with year, with site-specific variance components for species composition and plot). Abbreviations: df= nominator degree of freedom; ddf = denominator degree of freedom; logSR = log_2_(tree species richness); logShrubSR= log_2_(shrub species richness—this term is aliased with shrub presence and plot size and therefore fitted after these to only test for effects of shrub species richness in sub-plots of large plots where shrubs were present). *F* and *P* indicate F-ratios and P-values of the significance tests.

